# A Raman Spectroscopy-Based Method for Label-Free Discrimination of Human Inhibin *α*, Inhibin B, and Activin A

**DOI:** 10.64898/2026.07.04.735879

**Authors:** Weiyu Xiao, Yuzhen Dai, Sofia Martinez Gallardo Quijano, Domna Kotsifaki, Anastasia Tsigkou

## Abstract

Members of the transforming growth factor-β (TGF-β) superfamily, including inhibins and activins, are structurally related glycoprotein dimers that regulate reproductive and endocrine signaling. Their high degree of molecular similarity presents challenges for label-free analytical discrimination. To evaluate the ability of Raman spectroscopy to distinguish closely related TGF-β superfamily proteins based on intrinsic vibrational fingerprints. Raman spectra of recombinant human Inhibin *α*-subunit, Inhibin B β subunit, and Activin A (βA–βA) were acquired using confocal Raman microscopy with 532 nm excitation. Spectra were baseline-corrected, area-normalized, and analysed using principal component analysis (PCA). Distinct spectral signatures were observed across the 500–1800 cm^−1^ region. Differences within the S–S stretching region (500–550 cm^−1^) were consistent with variations in disulfide-bond environments, with the Inhibin *α*-subunit exhibiting the highest relative intensity in this region. Variations in the amide I band (1600–1700 cm^−1^) suggested differences in protein secondary structure, while aromatic amino acid vibrations provided additional discriminatory features. PCA revealed clear clustering and separation of all three protein classes based on their Raman fingerprints. Raman spectroscopy enables label-free differentiation of structurally related endocrine glycoproteins and demonstrates potential for the structural characterization and classification of inhibin and activin proteins within the TGF-β superfamily.

## 1. Introduction

The transforming growth factor-β (TGF-β) superfamily comprises more than 30 structurally related secreted cytokines that regulate embryogenesis, cell proliferation, differentiation, apoptosis, and endocrine homeostasis [1,2]. Within this family, inhibins and activins play central and opposing roles in reproductive endocrinology by regulating follicle-stimulating hormone (FSH) secretion from the anterior pituitary. Inhibins are heterodimeric glycoproteins composed of a common *α*-subunit linked via disulfide bonding to either a βA or βB subunit, forming inhibin A (*α*–βA) and inhibin B (*α*–βB). In contrast, activins are β-subunit dimers (βA–βA: activin A; βB–βB: activin B; βA–βB: activin AB) that stimulate FSH synthesis and secretion [3,4].

This functional antagonism is essential for hypothalamic–pituitary–gonadal axis regulation and folliculogenesis [4]. Beyond reproduction, dysregulated inhibin/activin signaling contributes to inflammation, fibrosis, tissue remodeling, and tumorigenesis in multiple organs, including ovary, prostate, and breast [5,6,7].

Structurally, both inhibins and activins are defined by the cystine-knot motif, consisting of a conserved disulfide-stabilized core and β-strand “finger-like” extensions [8,9,2]. However, incorporation of the inhibin *α*-subunit fundamentally alters both receptor specificity and biological function by enabling interaction with betaglycan, thereby converting activin-like signaling into antagonistic inhibin activity [10,11,3].

Despite high structural conservation among β-subunits, including 60–65% sequence identity and a shared cystine-knot fold, functional discrimination of inhibin and activin complexes remains analytically challenging. Conventional immunoassays often suffer from cross-reactivity and limited ability to distinguish free subunits from fully assembled dimers, while mass spectrometry typically requires extensive processing and does not preserve native conformations [12,8,3].

Raman spectroscopy (RS) provides a label-free alternative for probing protein structure by measuring vibrational modes following Raman scattering of monochromatic light. It is particularly sensitive to disulfide bonds (~500–550 cm^−1^), aromatic residues, and backbone amide I–III vibrations, which collectively report on secondary structure and higher-order assembly [13,14,15,16]. These spectral features are strongly influenced by hydrogen bonding, side-chain packing, and conformational constraints [17,18].

The cystine-knot scaffold is conserved across the TGF-β superfamily [19,20], yet ligands differ in disulfide topology, dimer interface geometry, and conformational flexibility [21]. These structural differences are expected to manifest as measurable variations in Raman spectral signatures, particularly within disulfide, aromatic, and amide regions.

In this study, we perform a comparative Raman spectroscopic analysis of recombinant Inhibin *α*-subunit, Inhibin B β subunit, and Activin A (βA–βA). Each protein exhibits a distinct vibrational fingerprint reflecting differences in cysteine network organization, aromatic residue environment, and quaternary structure. The isolated *α*-subunit is dominated by disulfide-associated vibrational modes consistent with a cysteine-rich scaffold, Inhibin B displays enhanced aromatic contributions consistent with interface stabilization, and Activin A shows broader amide and aromatic distributions indicative of increased conformational flexibility.

Collectively, these results demonstrate that Raman spectroscopy enables label-free discrimination of structurally homologous TGF-β family members by resolving intrinsic vibrational differences arising from disulfide geometry, aromatic packing, and dimerization state. This provides molecular-level structural discrimination that is not accessible through conventional immunochemical assays. From a translational perspective, this approach addresses a key limitation of ELISA and Western blot methodologies, which measure bulk immunoreactivity rather than structurally resolved bioactive states, limiting mechanistic interpretation in endocrine signaling [22,23]. By preserving native conformational information, Raman spectroscopy enables direct interrogation of structure–function relationships in a label-free manner, reframing diagnostic readouts as spectrally defined molecular states rather than concentration-based measures.

Raman spectroscopy provides direct molecular information that is sensitive to protein conformation and chemical composition [24]. This complementary capability is particularly relevant for endocrine disorders in which subtle shifts in *α*- and β-subunit balance may drive disproportionate biological effects. Resolving these structural states spectroscopically may therefore improve diagnostic specificity and mechanistic interpretation in TGF-β–related pathologies [12,22,23].

### 2. Results

### 2.1. Raman Spectral Profiling of TGF-β Superfamily Proteins

Raman spectra of recombinant human Inhibin *α*-subunit, Inhibin B β subunit, and Activin A (βA–βA) were recorded under identical acquisition parameters to ensure direct spectral comparability. Measurements were performed on solid-state protein preparations to preserve intrinsic vibrational modes associated with covalent bonding, disulfide architecture, and higher-order structural organization.

All three proteins exhibited characteristic Raman bands within the 500–1800 cm^−1^ fingerprint region, encompassing backbone, side-chain, and disulfide-associated vibrational modes. Despite their shared evolutionary origin within the TGF-β superfamily, each protein generated a distinct spectral fingerprint, indicating sensitivity of Raman spectroscopy to subunit composition and quaternary assembly [32,16,13].

### 2.2. Disulfide Bond Architecture as a Primary Spectral Discriminator

The disulfide (S–S) stretching region (500–550 cm^−1^) and C–S region (600–750 cm^−1^) provided the most prominent discriminating features. Inhibin *α* displayed the most intense S–S bands, consistent with its dense intramolecular disulfide network required for structural stabilization [30,33]. Activin A exhibited broader and less intense S–S features, consistent with a β–β homodimer architecture with distinct cysteine pairing topology. Inhibin B β subunit showed an intermediate spectral profile, reflecting its heterodimeric assembly and hybrid disulfide environment derived from *α* and βB subunits. These differences indicate that Raman spectroscopy is highly sensitive to cysteine connectivity and disulfide bond geometry, enabling discrimination of closely related glycoprotein assemblies [30,33].

### 2.3. Secondary Structure Contributions in the Amide I Region

The amide I band (1600–1700 cm^−1^), arising predominantly from C=O stretching vibrations of the peptide backbone, revealed subtle but reproducible shifts across the three proteins. Inhibin *α* exhibited a comparatively sharper amide I profile, consistent with a higher degree of ordered secondary structure [15,13].

Activin A displayed a broader amide I envelope, suggesting increased conformational flexibility and a higher relative contribution of β-sheet-like or disordered structural elements. Inhibin B exhibited composite spectral characteristics, reflecting its mixed *α*–β structural composition. These observations confirm that Raman spectroscopy can resolve secondary structure heterogeneity arising from subunit assembly differences.

### 2.4. Aromatic Residue Microenvironmental Sensitivity

Aromatic vibrations (1000–1600 cm^−1^) varied in intensity and band shape across the three proteins. Tyrosine and tryptophan modes showed shifted intensity ratios, indicating differences in hydrogen bonding and solvent exposure [13,33]. The phenylalanine ring breathing mode (~1004 cm^−1^) remained conserved across all samples, serving as a stable internal spectral reference. Variations in aromatic residue signatures likely reflect conformational constraints imposed by dimerization and glycoprotein folding.

Raman spectroscopy enables robust, label-free discrimination of structurally homologous TGF-β superfamily proteins by resolving differences in disulfide bonding, secondary structure, and aromatic microenvironment.

To establish a comparative baseline, Raman spectra of recombinant Inhibin *α*-subunit, Inhibin B β subunit, and Activin A (βA–βA homodimer) were acquired under identical experimental conditions. This allows direct evaluation of structural differences within the fingerprint region arising from subunit composition and quaternary assembly. The resulting spectra provide the foundation for subsequent structural interpretation across disulfide, backbone, and aromatic vibrational modes.

To first define the structural signature of an individual inhibin subunit, the Raman spectrum and molecular representation of the Inhibin *α*-subunit are presented. As a cysteine-rich monomer, this subunit is stabilized by an extensive intramolecular disulfide network, which is expected to strongly influence its vibrational fingerprint. Figure 1 highlights these structural features alongside its corresponding Raman spectral profile.

**Figure 1.**
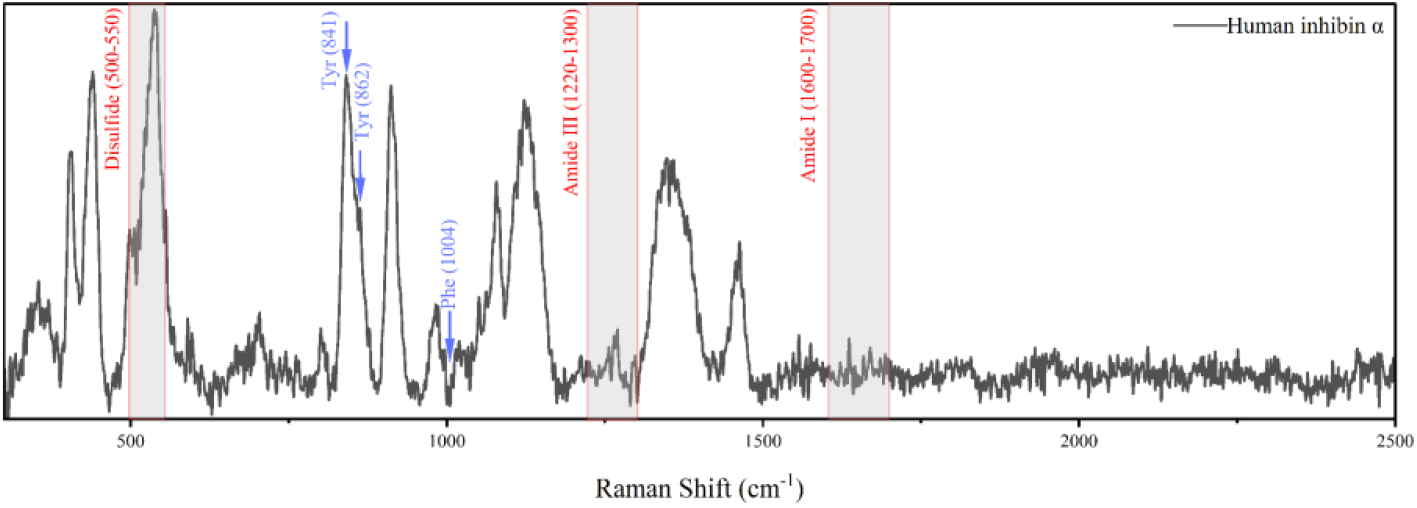
Raman spectrum of recombinant human inhibin *α*-chain. Baseline-corrected Raman spectrum of recombinant human inhibin *α*-subunit showing the principal vibrational bands associated with disulfide bonds, aromatic amino acid residues, and protein secondary structure. Major Raman bands corresponding to disulfide region (500–550 cm^−1^), tyrosine doublet (830/850 cm^−1^), phenylalanine (~1004 cm^−1^), amide III region (1220–1300 cm^−1^), and amide I region (1600–1700 cm^−1^) vibrations are annotated to facilitate interpretation of the molecular fingerprint.

**Figure 2.**
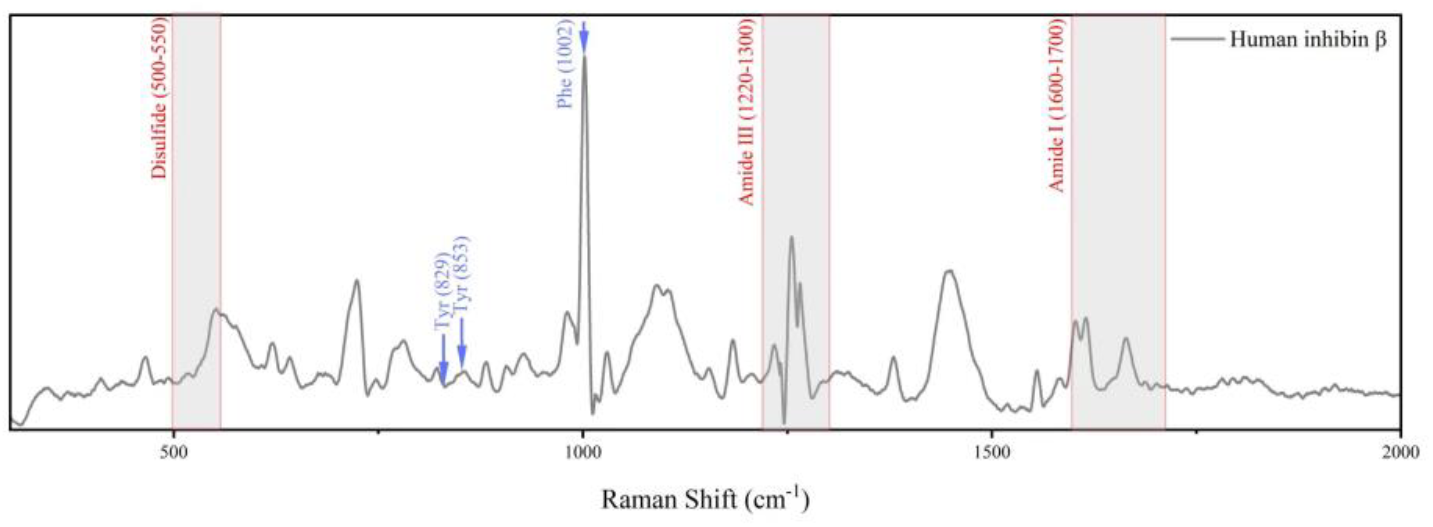
Raman spectrum of recombinant human Inhibin B acquired under identical experimental conditions. Spectrum highlights vibrational contributions in the disulfide region (500–550 cm^−1^), tyrosine doublet (830/850 cm^−1^), phenylalanine ring-breathing mode (~1004 cm^−1^), amide III region (1220–1300 cm^−1^), and amide I region (1600–1700 cm^−1^).

**Figure 3.**
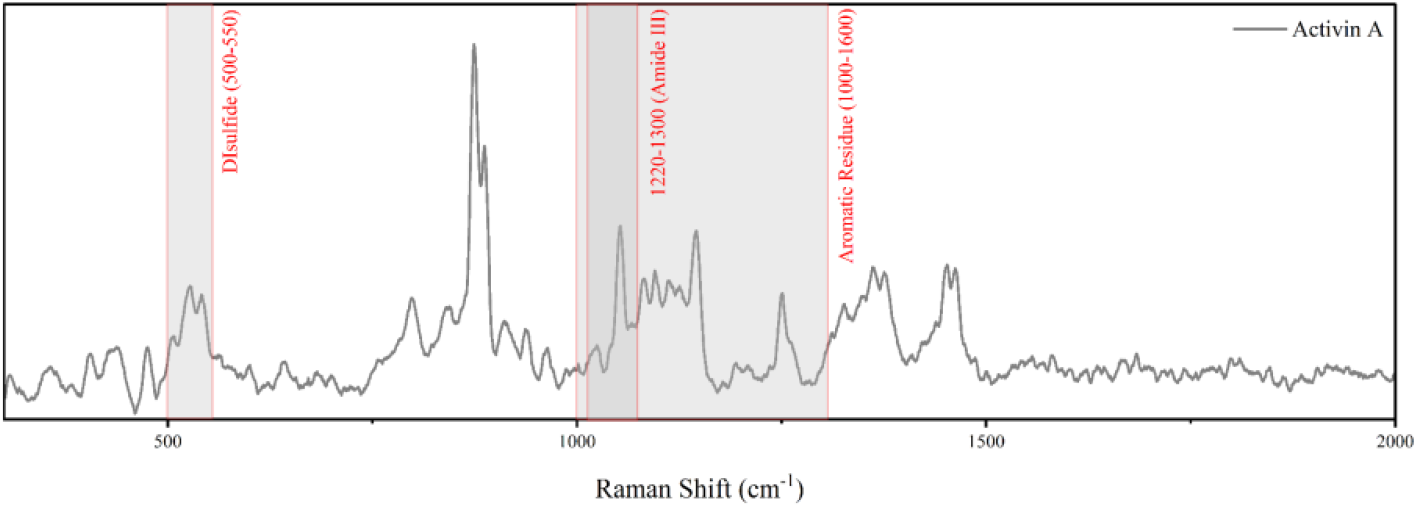
Raman spectrum of recombinant human Activin A measured under identical conditions. The spectrum shows characteristic bands in the disulfide region (500–550 cm^−1^), aromatic residue region (1000–1600 cm^−1^), and amide III region (1200–1300 cm^−1^).

**Figure 4.**
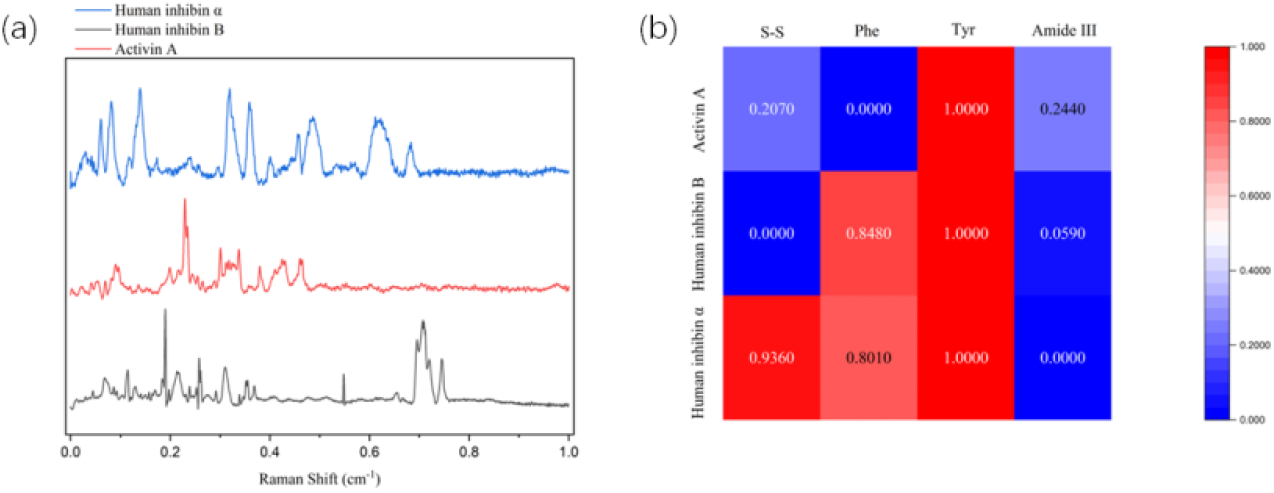
Comparative Raman fingerprints of recombinant human inhibin *α*, inhibin B, and activin A. (a) Baseline-corrected and normalized Raman spectra of recombinant human inhibin *α*-subunit (blue), inhibin B (black), and activin A (red), vertically offset for clarity. These proteins were selected because they are structurally related members of the transforming growth factor-β (TGF-β) superfamily that share a conserved cystine-knot architecture but differ in subunit composition and biological function. The comparison demonstrates the ability of label-free Raman spectroscopy to distinguish closely homologous proteins through characteristic differences in spectral features associated with disulfide bond conformations, aromatic amino acid residues, and protein secondary structure. (b) Heatmap showing normalized relative intensities of representative Raman bands corresponding to disulfide stretching (S–S, 500–550 cm^−1^), phenylalanine (Phe, ~1004 cm^−1^), tyrosine (Tyr, ~830–850 cm^−1^), and amide III (1200–1300 cm^−1^). The distinct spectral profiles highlight molecular differences between the inhibin *α*-subunit, heterodimeric inhibin B (*α*–βB), and homodimeric activin A (βA–βA), supporting Raman spectroscopy as a sensitive, label-free approach for discriminating structurally homologous proteins.

To examine the effect of heterodimerization on spectral features, Inhibin B (*α*–βB) is analyzed as a composite system comprising both *α* and βB subunits. This assembly introduces a mixed disulfide environment and altered secondary structure constraints relative to the isolated *α*-subunit. The following figure illustrates how these structural differences are reflected in its Raman fingerprint.

To complete the comparative framework, Activin A (βA–βA homodimer) is evaluated as a symmetric dimeric system lacking the *α*-subunit contribution. This configuration results in a distinct cysteine pairing topology and altered structural rigidity relative to both Inhibin *α* and Inhibin B. The following figure presents its Raman spectral signature in the context of these structural differences.

To establish a comparative structural framework across the inhibin–activin system, Raman spectra of Inhibin *α*-subunit, Inhibin B β subunit, and Activin A (βA–βA homodimer) were analyzed under identical experimental conditions. This enables direct assessment of how subunit composition and dimerization state influence vibrational signatures within the 500–1800 cm^−1^ fingerprint region. The comparative analysis highlights systematic differences in disulfide-associated modes, aromatic residue vibrations, and amide backbone features, reflecting progressive structural remodeling from monomeric to heterodimeric and homodimeric assemblies.

## 3. Discussion

The present study demonstrates that Raman spectroscopy can resolve subtle yet reproducible structural differences among closely related members of the TGF-β superfamily despite their extensive sequence homology and conserved cystine-knot architecture [1–4,20]. Using recombinant inhibin *α*, inhibin B (*α*–βB), and activin A (βA– βA), we identified unique vibrational fingerprints associated with disulfide bond organization, aromatic residue packing, and protein conformational dynamics [13–18,25,27].

These findings establish Raman spectroscopy as a sensitive, label-free approach for molecular discrimination of endocrine signaling proteins that are otherwise difficult to distinguish using conventional immunochemical techniques [22–25].

The transforming growth factor-β (TGF-β) superfamily includes numerous structurally homologous ligands that share high sequence identity and conserved cystine-knot folds yet elicit diverse and sometimes opposing biological functions [1,2,20]. Among these, inhibins and activins are paradigmatic examples: activins are β-subunit homodimers or heterodimers that signal through type I and type II serine-threonine kinase receptors, whereas inhibins incorporate a unique *α*-subunit that antagonizes activin signaling through betaglycan-dependent receptor modulation [10,11,26]. The ability to distinguish these proteins without labels or affinity reagents has remained a significant analytical challenge, particularly because antibodies frequently exhibit cross-reactivity toward shared β-subunit epitopes [3,4,12].

### 3.1. The Inhibin *α*-Subunit Exhibits a Unique Disulfide-Dominated Spectral Signature

The enhanced Raman intensity observed in the 500–550 cm^−1^ region for the inhibin *α*-subunit corresponds to S–S stretching vibrations associated with disulfide bonds [15– 18,30,31]. This observation is consistent with the highly cysteine-rich architecture of the inhibin *α*-subunit and the extensive intramolecular disulfide network required to stabilize the conserved cystine-knot motif characteristic of TGF-β family proteins [8,20].

Raman S–S stretching frequencies are known to be highly sensitive to disulfide bond geometry, torsional angles, and local conformational constraints [15–18,29,30]. Therefore, the strong disulfide-associated signal observed here likely reflects both the abundance and structural organization of disulfide bridges within the *α*-subunit scaffold. Biologically, this observation is significant because the *α*-subunit defines all inhibin isoforms and is absent from activins [3,4]. Incorporation of the *α*-subunit into β-subunit dimers fundamentally alters receptor interactions and downstream signaling pathways, converting activin-like signaling molecules into inhibin antagonists [10,11]. The preservation of a strong disulfide signature further suggests that the recombinant *α*-subunit retained its native tertiary architecture under the experimental conditions employed.

### 3.2. Inhibin B Displays Ordered Aromatic Packing

A prominent and highly resolved phenylalanine ring-breathing mode near 1003 cm^−1^ was observed in inhibin B β subunit. This vibration is a well-established Raman marker of aromatic side-chain organization and local protein packing [13,15,27,32]. The intensity and sharpness of this feature indicate that phenylalanine residues within the inhibin B β subunit occupy highly ordered environments characterized by restricted rotational freedom and reduced conformational heterogeneity.

Such behavior is consistent with structural studies demonstrating that dimerization within members of the TGF-β superfamily generates defined intermolecular interfaces that constrain aromatic side-chain orientation [8,20]. In contrast, the corresponding aromatic bands in activin A were broader and less intense, suggesting increased conformational flexibility.

The observed differences cannot be attributed solely to amino acid composition because both proteins contain aromatic residues. Rather, they likely reflect distinct quaternary structural organizations and differences in dimer-interface packing [8,20]. Consequently, the phenylalanine marker band may serve as a useful spectroscopic indicator of heterodimer assembly and structural integrity.

### 3.3. Amide III Broadening Suggests Greater Conformational Plasticity in Activin A

The amide III region (1200–1300 cm^−1^) and amide I region (~1655 cm^−1^) provide important information regarding protein secondary structure and conformational heterogeneity [13–18,25]. Activin A exhibited broader spectral envelopes across these regions than either inhibin *α* or inhibin B β subunit.

Raman band broadening is commonly associated with increased conformational flexibility, dynamic disorder, or the coexistence of multiple structural substates [14,17,18]. Accordingly, the broader amide bands observed in activin A may indicate a more flexible βA–βA homodimeric architecture relative to the more constrained inhibin B β subunit.

This conformational plasticity may have functional implications. Activins interact with multiple receptor combinations and participate in a wide range of biological processes, including reproductive regulation, inflammation, tissue repair, and cellular differentiation [4,6,21,26]. Increased structural flexibility may facilitate receptor recognition and signaling versatility, whereas the more rigid architecture of inhibins may support specific receptor-modulatory functions mediated through betaglycan [10,11].

### 3.4. Principal Component Analysis Confirms Statistical Discrimination

Principal component analysis demonstrated clear separation of inhibin *α*, inhibin B β subunit, and activin A spectra, confirming that the observed spectral differences are sufficiently robust for statistical classification. Separation was driven primarily by variations in disulfide-associated bands, aromatic residue signatures, and amide backbone vibrations [13–18,25].

These findings indicate that Raman spectroscopy captures higher-order structural information rather than merely reflecting differences in protein abundance. This capability is particularly important because many conventional immunoassays measure bulk immunoreactivity without providing direct information regarding protein folding state, dimerization status, or conformational integrity [12,22].

### 3.5. Translational Implications and Future Directions

From a translational perspective, the ability to distinguish TGF-β family proteins without labels or affinity reagents opens new opportunities for spectroscopic biosensing and structural diagnostics [22–24]. Altered inhibin and activin signaling has been implicated in infertility, polycystic ovary syndrome, endometriosis, ovarian cancer, breast cancer, fibrosis, and cardiovascular disease [5–7,12,26,34].

Current clinical measurements rely primarily on ELISA-based assays that provide quantitative information but limited structural insight [12]. Raman spectroscopy offers a complementary analytical approach capable of simultaneously providing compositional and conformational information in a label-free format [22–25].

Integration with portable Raman instrumentation, surface-enhanced Raman spectroscopy (SERS), and machine-learning-assisted spectral classification may ultimately facilitate rapid point-of-care assessment of reproductive and endocrine biomarkers [22,23].

### 3.6. Limitations

Several limitations should be acknowledged. First, measurements were performed using purified recombinant proteins under controlled experimental conditions and therefore may not fully recapitulate the complexity of biological fluids. Second, Raman spectroscopy provides indirect structural information and should ideally be complemented by orthogonal approaches such as circular dichroism spectroscopy, X-ray crystallography, cryo-electron microscopy, and molecular dynamics simulations [8,20,21,31].

Finally, additional members of the TGF-β superfamily, including inhibin A, activin B, activin AB, BMPs, GDFs, and TGF-β isoforms, should be examined to establish the broader applicability of the approach. Future studies incorporating larger spectral datasets and machine-learning classification frameworks may enable automated identification of TGF-β family proteins with high diagnostic accuracy [22,23].

## 4. Materials and Methods

All experiments were performed under controlled conditions, with temperature maintained at 25 °C and relative humidity monitored and stabilized within the chamber to prevent spectral artifacts.

### 4.1. Protein Samples

Recombinant human inhibin *α*-subunit (INHA; Abcam, Cambridge, UK; Cat. No. ab280388), recombinant human inhibin B β subunit (INHBB; Abcam, Cambridge, UK; Cat. No. ab158764), and recombinant human activin A (βA–βA homodimer; R&D Systems, Minneapolis, MN, USA; Cat. No. 11344-AC) were obtained from commercial suppliers and used according to manufacturer specifications. All proteins were stored under recommended conditions until analysis.

### 4.2. Sample Preparation

To preserve the native molecular architecture of the recombinant proteins and avoid potential conformational alterations introduced by reconstitution, all samples were analyzed directly in their commercially supplied lyophilized state. Inhibin *α* (INHA), inhibin B β subunit, and activin A were examined using Raman spectroscopy. Approximately 2 μg of each lyophilized protein was deposited onto a glass microscope slide and analyzed without further processing. All measurements were performed using identical optical, instrumental, and spectral acquisition parameters to facilitate rigorous inter-protein comparison. For each protein, two independent sample preparations were evaluated, with Raman spectra acquired from 3–5 spatially distinct, non-overlapping locations per preparation to assess spatial uniformity and experimental reproducibility. High spectral agreement between replicate measurements confirmed the reliability of the solid-state Raman fingerprinting approach.

### 4.3. Raman Spectral Acquisition

Raman spectra were acquired using a confocal Raman microscope (Horiba Scientific, Kyoto, Japan) equipped with a motorized XYZ stage for automated spatial mapping. Excitation was provided by a 532 nm continuous-wave diode laser focused onto the sample.Laser power at the sample plane was maintained below 15 mW to prevent local heating or photodegradation. Rayleigh scattering was suppressed using a notch filter, and Raman-scattered photons were dispersed using an 1800 lines/mm grating and detected with a thermoelectrically cooled CMOS detector. The spectrometer was calibrated prior to each measurement session using the silicon standard band at 520.7 cm^−1^. Spectra were collected over the range 300–2800 cm^−1^, with primary analysis focused on the fingerprint region (300–2000 cm^−1^). Acquisition parameters were set to 15 s integration time with two accumulations per spectrum. Background spectra acquired from clean glass substrates under identical conditions were subtracted from all measurements.

### 4.4. Spectral Preprocessing and Peak Assignment

Raw Raman spectra were processed using OriginPro 2025b (OriginLab Corporation, Northampton, MA, USA). Spectra exhibiting excessive noise, cosmic ray artifacts, or acquisition instability were excluded from analysis.Fluorescence background was corrected using polynomial baseline subtraction. All spectra were subsequently normalized using total area normalization over the full spectral range (300–2800 cm^−1^), enabling intensity-independent comparison across samples.

Peak assignments were performed based on established Raman protein spectroscopy databases and literature reports, with emphasis on disulfide bond vibrations, amide modes, and aromatic residue contributions (Table 1) [13–18,25,27–31].

**Table 1.** Peak assignments were made based on established Raman spectroscopic literature for proteins and biomolecules, encompassing general vibrational modes of proteins [13,14,27], aromatic amino acid marker bands [28,29], and disulfide-rich glycoprotein structural studies relevant to cystine-knot growth factors of the TGF-β superfamily [30,31].

| Raman shift (cm<sup>-1</sup>) | Assignment | Vibrational mode | Key references |
| --- | --- | --- | --- |
| 500-550 | S-S stretching | Disulfide bond stretching | [15,17,18] |
| 830/850 | Tyrosine residue | Ring breathing coupled + overtone vibration | [28,15,18] |
| ~1004 | Phenylalanine ring breathing | Aromatic ring breathing | [15,13,25] |
| 1220–1300 | Amide III | C–N stretch + N–H bend | [15,17,25] |
| 1600-1700 | Amide I / C=C | Predominantly C=O stretching of peptide backbone; aromatic C=C stretching may contribute | [15,17,25] |

### 4.5. Statistical Analysis

Principal component analysis (PCA) was performed using OriginPro 2025b (OriginLab Corporation, Northampton, MA, USA). All spectra were baseline-corrected, area-normalized, and mean-centered prior to PCA. Hierarchical clustering and heatmap visualization were generated using the same software package.

## Author Contributions

Investigation, methodology, formal analysis and visualization (figures), W.X. and Y.D.; writing of the original draft, W.X. and Y.D.; contribution to the introduction and the references, review and editing, proofreading and template preparation, S.M.G.Q.; conceptualization, supervision and project administration, D.G.K. and A.T. All authors have read and agreed to the published version of the manuscript.

## Institutional Review Board Statement

Not applicable. This study used only commercially obtained recombinant proteins and did not involve humans or animals.

## Informed Consent Statement

Not applicable.

## Data Availability Statement

Suggested Data Availability Statements are available in the section “MDPI Research Data Policies” at https://www.mdpi.com/ethics.

## Acknowledgments

We would like to sincerely thank Professor Chengcheng Zheng and the Optics Laboratory for generously allowing us to use their facilities at Duke Kunshan University Division of Natural and Applied Sciences. We are also deeply grateful to Yuli for her invaluable training and technical support throughout the project. The authors acknowledge the use of generative AI tools for language editing and manuscript polishing. All scientific content, interpretations, and conclusions were independently developed and verified by the authors.

